# Identification of enzymes for lignocellulose degradation in the genome and transcriptome of the aquatic hyphomycete *Clavariopsis aquatica*

**DOI:** 10.1101/2020.06.18.151886

**Authors:** Felix Heeger, Elizabeth C. Bourne, Christian Wurzbacher, Elisabeth Funke, Anna Lipzen, Guifen He, Vivian Ng, Igor V. Grigoriev, Dietmar Schlosser, Michael T. Monaghan

## Abstract

Fungi are ecologically important decomposers of lignocellulose. Basidiomycetes use peroxidases, laccases, and enzymes of the cytochrome P450 superfamily for cometabolic lignin degradation in order to access cellulose and hemicellulose as carbon sources. Limited lignin modification capabilities have also been reported for some terrestrial ascomycetes. Here we newly sequenced the genome of an exclusively aquatic ascomycete, *Clavariopsis aquatica*, documented the presence of genes for the modification of lignocellulose and its constituents, and compared differential gene expression between *C. aquatica* cultivated on lignocellulosic and sugar-rich substrates. We identified potential peroxidases, laccases, and cytochrome P450 monooxygenases several of which were differentially expressed when experimentally grown on different substrates. Additionally, we found regulation of pathways for cellulose and hemicellulose degradation. Our results suggest that *C. aquatica* is able to modify lignin, detoxify aromatic lignin constituents, or both. This may facilitate the use of carbohydrate components of lignocellulose as carbon and energy sources.

## Introduction

Fungi are important decomposers of lignocellulose, which is the main component of plant cell walls and contains cellulose, hemicellulose, and lignin. Lignin is the most recalcitrant among these polymers and is composed of phenylpropanoid monomer units. The proportions of the three components of lignocellulose vary among plant materials and plant species. Lignocellulose with the highest proportion of lignin occurs in wood, with wood decay predominantly mediated by fungi from the phylum Basidiomycota. Of these, only the so-called “white rot” fungi are able to completely mineralize lignin, while “brown rot” fungi can only modify it to some extent. White rot fungi express multiple groups of extracellular lignin-modifying enzymes such as laccases and various peroxidases (Lundell et al., 2010). Comparative studies of the genomes of wood-decaying white and brown rot fungi have provided insights into the protein families that are important for lignocellulose degradation (Floudas et al., 2012; Frommhagen et al., 2017; Riley et al., 2014). These studies also concluded that white and brown rot fungi are neither monophyletic nor easily distinguishable by their protein repertoire. Greater insight into the proteins that are produced during wood degradation has been gained by studies of gene expression (e.g. Tang et al., 2013; Yang et al., 2012). Ries et al. (2013) studied terrestrial ascomycetes and found that, like basidiomycotes, they also possess and express genes for lignocellulose degradation, where cellulose and hemicellulose components are the preferred targets.

In plant material that is less lignin-rich than wood, other fungi can break down cell walls and play an important role in degradation. One such example is leaf litter that becomes submerged in streams, where aquatic hyphomycetes of the phylum Ascomycota are often the most abundant fungal species (Duarte et al., 2015). It is generally accepted that aquatic hyphomycetes can degrade cellulose and hemicellulose from plant litter, but their ability to degrade lignin is less well studied (Gessner et al., 2007; Krauss et al., 2011).

*Clavariopsis aquatica* is an exclusively aquatic ascomycete (aquatic hyphomycete) that effectively colonizes leaf litter in streams (Iqbal and Webster, 1973; Suberkropp and Klug, 1976). It has been reported to biotransform environmental pollutants such as nonylphenol and polycyclic musk fragrances in a cometabolic manner, thereby involving both extracellular laccase and intracellular oxidation reactions that are indicative of the action of cytochrome P450 systems (Junghanns et al., 2005; Krauss et al., 2011; Martin et al., 2007). In basidiomycetes, both enzyme classes have been implicated in the oxidation of plant-derived compounds such as lignin constituents of lignocellulose (Floudas et al., 2012; Lundell et al., 2010; Riley et al., 2014; Syed et al., 2014). Because of the various and sometimes multiple functions of these protein families, and the frequent occurrence of their members in many different forms, the function of the identified genes cannot easily be inferred from their sequence alone.

*C. aquatica* represents a model organism for the ecological group of aquatic hyphomycetes, also known as freshwater hyphomycetes or Ingoldian fungi (Krauss et al., 2011). Its ability to effectively degrade plant material and xenobiotics under submerged conditions render it potentially interesting for biotechnological applications; however, the molecular basis of plant decomposition by aquatic hyphomycetes has not yet been investigated in detail.

In this study, we newly sequenced the genome of *C. aquatica* and searched for laccase and cytochrome oxidase enzyme systems, and for peroxidases known to act on lignocellulose components in other fungi. We also used a replicated incubation assay to examine differential gene expression during growth on substrates with varying lignin content (alder, wheat, malt). Common alder (*Alnus glutinosa*) leaves are a typical natural substrate of the *C. aquatica* in rivers. Wheat straw is a substrate that is not typically found in *C. aquatica* habitats, but has a higher cellulose content and higher proportion of lignin than alder leaves (Alemdar and Sain, 2008; Bjerre et al., 1996; Chauvet, 1987; Lecerf and Chauvet, 2008), and thus might prompt a stronger signal of differential expression of genes related to the metabolism of these polymers. Malt extract is a mainly sugar-based substrate, devoid of phenolic and other aromatic constituents, and served as a control. We examined annotations of differentially expressed genes for over-represented annotation terms in an effort to identify critical fungal pathways that may be involved in fungal carbon cycling under submerged conditions. To our knowledge, this is the first combined genome and gene expression study of an exclusively aquatic fungus.

## Methods

### Cultivation

Liquid cultivations of *C. aquatica* were carried out in 500-mL flasks containing 200 mL of medium. For cultivation, 10 g/L milled alder leaves or wheat straw (particle size ca. 2-4 mm), were autoclaved (121 °C, 20 min) twice and suspended in a nitrogen-limited solution previously described for manganese peroxidase production in *Stropharia rugosoannulata* (Schlosser and Höfer, 2002). Control cultures were grown on liquid malt extract medium (1% malt extract, w/v; pH 5.6-5.8) (Solé et al., 2012). The flasks were inoculated with 5 mL of a mycelial suspension of *C. aquatica* prepared in sterile water (Junghanns et al., 2005). Cultures were agitated at 120 rpm and incubated at 14 °C in the dark. Flasks were harvested after 7 days (trophophase/exponential growth phase) and 20 days (stationary growth phase) of cultivation (Junghanns et al., 2005; Solé et al., 2012). Liquid media were removed by stepwise centrifugation of the content of a flask in sterile 50-mL conical tubes at 7197 g and 4 °C for 10 min (Eppendorf centrifuge 5430R, rotor FA-35-6-30; Eppendorf, Hamburg, Germany). After discarding respective supernatants, biomass pellets obtained from one cultivation flask were combined in one sterile 50-mL conical tube, shock frozen in liquid nitrogen, and kept frozen at −80 °C until RNA extraction.

Fungal growth phases and related starvation conditions are well known to play important roles in the regulation of enzymes involved in lignocellulose degradation (Delmas et al., 2012; Solé et al., 2012). Liquid culturing with milled plant material allows for clear temporal differentiation between exponential and stationary growth phase, and was applied in this study. To investigate more natural conditions, additional cultures were grown on comparatively more inhomogeneous solid substrate (i.e. not suspended in solution/water) where different fungal growth phases may potentially co-exist at the same time point.

For cultivation on solid substrate (wheat straw or alder leaves), 100-mL flasks were supplemented with 2 g (dry mass) of milled substrate (about 2-4 mm particle size) and 8 mL of tap water, and autoclaved (121 °C, 20 min) twice. The flasks were inoculated with six mycelia-containing agar plugs, derived from the edge of *C. aquatica* colonies on malt agar plates (Junghanns et al., 2005), and incubated without agitation at 14 °C in the dark. Flasks were harvested after 26 days of cultivation, solid substrates were shock frozen in liquid nitrogen, and kept frozen at −80 °C until RNA extraction.

Overall, there were three conditions (liquid culture during stationary growth, liquid culture during exponential growth, and solid-state culture) for each of the natural substrates (wheat straw, alder leaves) and two conditions (liquid culture during stationary and exponential growth phases) for malt extract. All sampling was performed in triplicate, which led to 24 samples in total. Sterile conditions were ensured throughout sampling.

### Genome Sequencing

*C. aquatic*a WD(A)-00-1 mycelium was extracted from agar plates and subjected to DNA extraction using the FastDNA Spin Kit for Soil (MP Biomedicals, USA) following the manufacturer‘s instructions. Whole-genome shotgun reads of the *C. aquatica* genomic DNA were generated using a NexteraXT library preparation kit (Illumina, USA) following the manufacturer’s protocol and sequenced on a MiSeq (paired-end, 300bp) instrument with the v3 chemistry (Illumina) after library verification with a Nano Kit (Illumina).

### RNA-Seq

Frozen material from each sample was ground to a fine powder using an RNase-cleaned and pre-cooled pestle and mortar and liquid nitrogen, with a small spatula of zirconium beads (Biospec, USA) added for additional friction. RNA extraction followed Bourne et al. (in press) and Johnson et al. (2012) using a CTAB-based extraction. Briefly, for each sample, ca. 500 mg of ground, frozen tissue was added to 1.4 mL pre-heated (65 °C) CTAB buffer, vortexed until thoroughly mixed, incubated at 65 °C for 10-15 min, and centrifuged at 13,000 g for 3 min. The supernatant was transferred to a new 2-mL tube for two rounds of chloroform:isoamyl (24:1) extraction, a single phenol-chloroform extraction (5:1, pH 4.5), and a final chloroform:isoamyl (24:1) extraction. Following centrifugation, the upper phase was transferred to a new 2-mL tube. Purification was performed using the RNeasy Mini Kit (Qiagen, Germany) with on-column DNA digestion (RNase-free DNase set, Qiagen), following the manufacturer’s guidelines. RNA was eluted by adding 30 µL of elution buffer directly to the membrane and spinning at 13,000 g for 1 minute.

Total RNA was quantified using the QuantiFlour RNA system (Promega, USA). The presence of DNA was checked using the QuantiFlour DNA system (Promega), and samples with remaining DNA underwent an additional post-extraction DNase treatment using the TURBO DNA-*free* Kit (Invitrogen, Thermo Fisher Scientific, USA) following the manufacturer’s guidelines. Integrity of the RNA was assessed with the Agilent RNA 6000 Nano Kit and Agilent 2100 Bioanalyzer (Agilent Technologies, USA) following manufacturer’s guidelines. The RNA integrity (RIN) value was determined as a proxy of the overall quality of the RNA sample, with a value greater than 6 considered suitable for further analysis. We also assessed the quality of the overall Bioanalyzer trace by eye. Multiple extractions were performed for each sample and pooled to obtain sufficient RNA quantity for sequencing. Samples with sufficient quality were sent on dry ice for library preparation and sequencing (see below). RNA of sufficient quantity and quality could not be obtained from all samples. The sampling points for stationary growth in liquid culture and on the solid substrate for alder leaves had to be excluded, as well as for stationary growth on malt extract. For exponential growth in liquid culture for alder leaves, only two of the replicates produced suitable quantities of RNA. In total, 14 samples were used for library preparation.

RNA library preparation and sequencing was performed at the DOE Joint Genome Institute in Walnut Creek, CA, USA. Stranded cDNA libraries were generated using the Illumina Truseq Stranded RNA LT kit. mRNA was purified from 1 µg of total RNA using magnetic beads containing poly-T oligos. mRNA was fragmented and reverse-transcribed using random hexamers and SSII (Invitrogen), followed by second strand synthesis. The fragmented cDNA underwent end-repair, A-tailing, adapter ligation, and 8 cycles of PCR. qPCR was used to determine the concentration of the libraries. Libraries were sequenced on the Illumina Hiseq (single-end, 100bp).

### Genome Assembly

Reads were digitally normalized with khmer (version 0.7.1, Crusoe et al., 2015) to remove read errors and reduce computation times, as follows: In a first step, reads were normalized to a coverage of 20 (Brown et al., 2012). After removal of low-abundance kmers (Zhang et al., 2014), another round of normalization to a non-redundant coverage of 5 was applied (Supplemental Info 1) and only read pairs with both reads remaining after normalization were used for assembly. Assembly was performed with velvet (version 1.2.10, Zerbino and Birney, 2008) and run with different kmer lengths k (see Supplemental Info 1 for details). Finally, k=27 was chosen because it resulted in the highest N50 score. To estimate genome completeness, we ran BUSCO (version 3.0.2, Simão et al., 2015) with the pezizomycotina reference set of single-copy genes. The *clean* command of the funannotate pipeline (version 1.2.0, Jon Palmer and Jason Stajich, 2019) was used to remove contigs shorter than 500 bp as well as redundant contigs.

### Genome Annotation

Reads obtained from RNA-Seq were *de novo* assembled into putative transcripts with Trinity (version 2.5.1, Grabherr et al., 2011). Trinity was configured to use trimmomatic for trimming and to perform digital normalization (see Supplemental Info 1 for further details). Normalized RNA-Seq reads produced by Trinity were mapped to the genome contigs with Star (version 2.5.3a, Dobin et al., 2013), using default parameters. The mapped reads were used to generate a genome-guided assembly with Trinity (see Supplemental Info 1 further details). The PASA pipeline (version 2.2.0, Haas et al., 2003) was used to combine *de novo* and genome-guided assembled transcripts into a single gff file as evidence for annotation (Supplemental Info 1). The resulting gff file together with genome-guided assembled transcripts and mapped reads were used as input for the *predict* command of the funannotate pipeline. The *update* command of funannotate was then used to add UTR annotations. The predicted protein sequences were used as input for Interproscan (version 5.27, Jones et al., 2014) to generate Interpro (Finn et al., 2017) as well as Gene Ontology (GO, Ashburner et al., 2000; The Gene Ontology Consortium, 2017) annotations. The *annotate* command of the funannotate pipeline was used to combine Interproscan results with CAZy (Lombard et al., 2014) annotations from dbCAN (version 6.0, Yin et al., 2012).

In addition to the annotations from the funannotate pipeline (above), proteins were assigned as either secreted or not secreted using signalP (version 4.1, Petersen et al., 2011) and to KEGG (Kyoto Encyclopedia of Genes and Genoms, Kanehisa et al., 2016a) orthology groups with the BlastKOALA web service (Kanehisa et al., 2016b).

Besides general genome annotation, we specifically searched for gene families known to be involved in lignin degradation. We performed a blast search against our newly assembled *C. aquatica* genome to check for five previously described, partially sequenced laccase genes (Solé et al., 2012). In addition, we identified multicopper oxidases (MCO) by assignment to the CAZy family AA1. MCOs were further classified using a blast search of the Laccase and Multicopper Oxidase Engineering Database (version 6.4, Sirim et al., 2011). We identified possibly relevant peroxidases by annotation with the Interpro family IPR001621 (Fungal ligninase) and verified the resulting proteins by annotation with the Peroxiscan (Koua et al., 2009) web service (accessed May 15^th^ 2018) of PeroxiBase (Fawal et al., 2013). Proteins potentially belonging to the cytochrome P450 family were identified by annotation with the Interpro family IPR001128 (Cytochrome P450).

### Differential Expression and MGSA Analysis

Read counts per gene were generated with RSEM (version 1.3, Li and Dewey, 2011) using default parameters from mapped RNA-Seq reads (see above). All of the following analyses were implemented as a snakemake (version 3.5.4, Köster and Rahmann, 2012) workflow that can be found at www.github.com/f-heeger/caquatica_expression. RSEM output files were combined into a single read count matrix with the *merge_RSEM_output_to_matrix*.*pl* script from Trinity. Differential gene expression between different samples was modeled with the DESeq2 (version 1.10.1, Love et al., 2014) R package. Genes with an adjusted p-value < 0.05 and an absolute log2 fold change > 1 were considered to be differentially expressed.

Multiple Gene Set Activation (MGSA) analysis uses a Bayesian network approach to predict a probability of activation for sets of genes for each comparison (e.g., wheat straw – alder) based on differentially expressed genes (Bauer et al., 2011, 2010). We defined gene sets in three ways for the MGSA analysis using three different annotations: (1) all genes annotated with one GO term, (2) all genes assigned to one CAZy family, and (3) all genes assigned to one KEGG pathway based on assignment to KEGG orthology groups (see above). The activation probability cut-off, above which a gene set is considered to be “activated”, is ultimately arbitrary. The authors of the method suggest to use 0.5 (Bauer et al., 2010), reasoning that this means that the gene set is “more likely to be on than to be off”. We chose a slightly more conservative cutoff of 0.6 for gene sets to be considered activated. We note that activation is a statistical term, here indicating that differential expression of genes in these sets can be best explained by some form of regulation of these sets, given the Bayesian model underlying the MGSA analysis.

## Results

### Genome Assembly and Annotation

We obtained 7.31 million read pairs (2×300 bp), which were deposited in the NCBI Sequence Read Archive under the acession PRJNA610219. They were reduced to 1.09 million reads by digital normalization and assembled into 2,650 non-redundant contigs (longer than 500bp) with a N50 score of 30,079 bp and a total length of 34.18 Mb. These included complete single copies of 94.8% of the single-copy genes expected according to the BUSCO reference set, indicating good completeness of our assembly. The genome assembly was deposited at DDBJ/ENA/GenBank under the accession JAAZQE000000000 and can be found in Mycocosm (Grigoriev et al., 2014) under mycocosm.jgi.doe.gov/Claaq1. A total of 12,100 proteins were predicted by the funnanotate pipeline, of which 6,128 (50.64%) were annotated with at least one GO term, 2,322 (19.19%) were assigned to at least one KEGG pathway, and 572 (4.73%) to at least one CAZy family. There were 5,724 (47.31%) proteins that did not receive any annotation from these databases.

All five of the laccase gene sequences previously described from *C. aquatica* (Solé et al., 2012) were present in our genome, with nucleotide identity > 98%. Based on annotation with CAZy auxiliary activity family AA1, we identified all five known laccases and eight additional multicopper oxidases. They all exhibited a high degree of similarity (53-100% pairwise amino acid identity) for conserved sites of laccase genes (Kumar et al., 2003) in a multiple alignment (data not shown). When compared to the Laccase and Multicopper Oxidase Engineering Database with blast, only one of the previously described laccases (lcc2) was classified as belonging to the “Basidomycete Laccase” superfamily in the database. The other four were assigned to the superfamily “Ascomycete MCO”. Of the newly identified potential multicopper oxidases, five were assigned to “Ascomycete MCO” as well, while the other three were classified as “Fungal Ferroxidase” and will not be considered further.

Based on annotation with the Interpro family IPR001621, we identified five peroxidases. Annotation with Perociscan, verified all as belonging to the class II peroxidase superfamily in PeroxiBase and further assigned them to the “Asco Class II” family.

A total of 137 proteins were identified as belonging to the cytochrome P450 superfamily by annotation with the Interpro family IPR001128.

### RNA-Sequencing and Differential Expression

We obtained 317.18 million RNA-Seq single end reads (100 bp) in total, with > 14 million reads for each sample (see Table 1). Reads were deposited in the NCBI Sequence Read Archive under the accessions PRJNA440444 - PRJNA440457. Most (75.27%, SD 1.33%) of the reads from each sample could be mapped to our newly assembled *C. aquatica* genome with RSEM. Two of the samples (both liquid culture, exponential phase on wheat straw) had considerably more reads than the rest (50.50 and 57.49 million). Sub-sampling to 17 million reads (rounded mean number of reads in the other samples) and re-running RSEM mapping and differential expression analysis with DESeq2 showed only minor differences (97.55% genes with the same expression status). Because of this result and considering that read count per sample is accounted for in the DESeq2 model, we used all reads without sub-sampling for further analyses. We modeled differential expression between recalcitrant and nutrient-rich media (wheat straw versus malt extract and alder versus malt extract), between the two growth phases on wheat straw (stationary versus exponential), and among methods of culture (solid culture versus exponential growth in liquid culture and solid culture versus stationary growth in liquid culture).

**Table 1:**
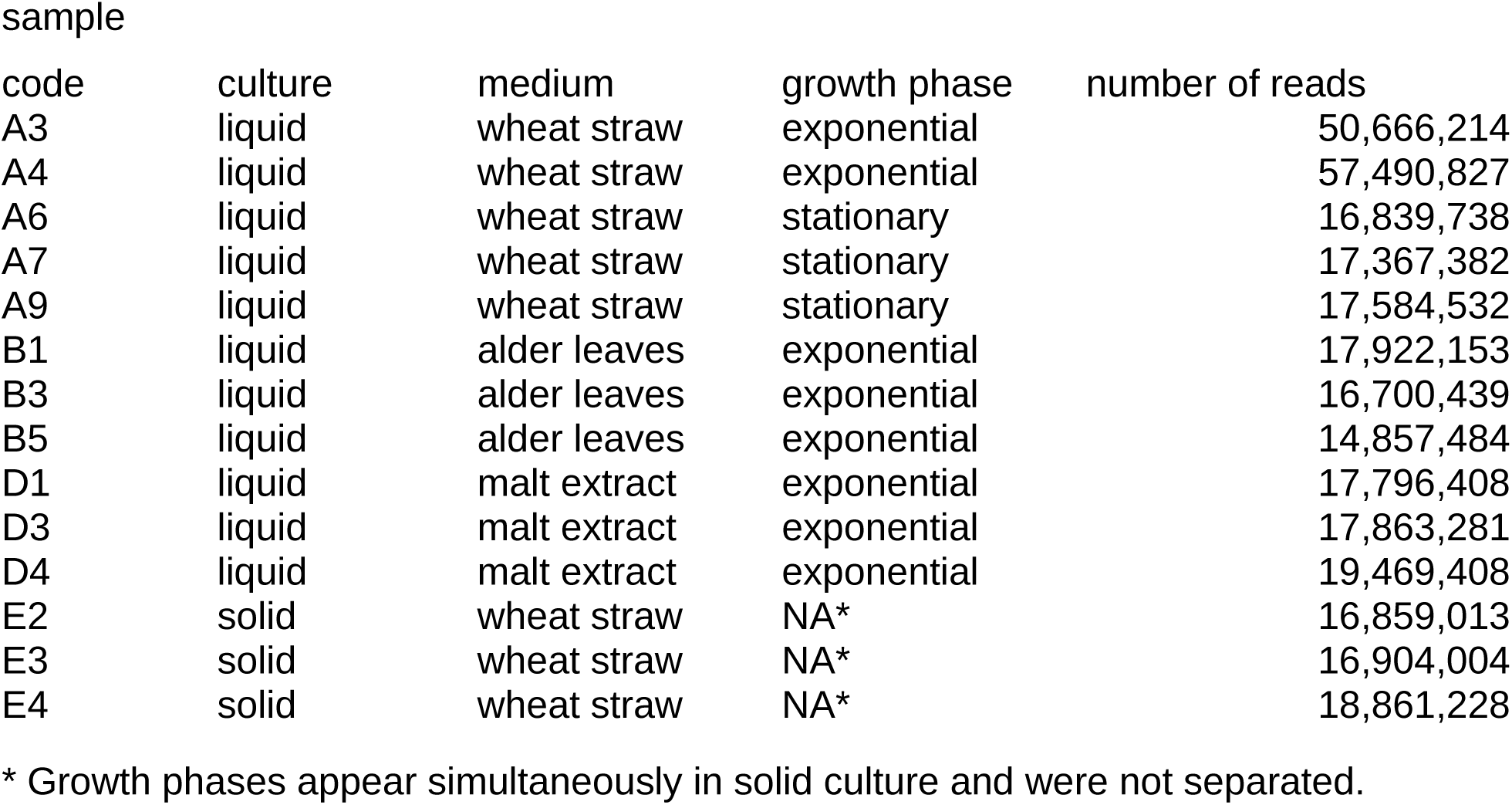
*C. aquatica* samples grown under different conditions.

Gene expression showed strong differences between solid and liquid cultures, with the most differentially expressed genes being between growth in solid culture compared to stationary growth in liquid culture, and the second-most differentially expressed genes between solid growth and exponential growth in liquid culture. The difference between stationary and exponential growth was weaker by comparison (see Table 2). Growth on both plant matter substrates induced similar changes in the metabolism compared to malt extract, indicated by a significant (Fisher’s exact test, p<10^−192^) overlap between differentially expressed genes.

**Table 2:** Number of up- and down-regulated genes for different comparisons

| comparison |  | differential expression |  |
| --- | --- | --- | --- |
|  |  | up-regulated | down-regulated |
| condition 1 | condition 2 |  |  |
| exponential growth on wheat | exponential growth on malt |  |  |
| straw in liquid culture | extract in liquid culture | 1,430 | 1,570 |
| exponential growth on alder | exponential growth on malt |  |  |
| leaves in liquid culture | extract in liquid culture | 1,033 | 1,462 |
| exponential growth on wheat | stationary growth on wheat |  |  |
| straw in liquid culture | straw in liquid culture | 1,380 | 883 |
| growth on wheat straw in solid | exponential growth on wheat |  |  |
| culture | straw in liquid culture | 2,731 | 2,328 |
| growth on wheat straw in solid | stationary growth on wheat |  |  |
| culture | straw in liquid culture | 2,683 | 2,478 |

The five known laccase genes, as well as the eight newly identified laccase-like genes, showed no consistent pattern of up- or down-regulation in the exponential growth phase in liquid culture when comparing alder and wheat straw (Fig. 1). Of the six identified putative Class II peroxidases, two were up-regulated on wheat straw and one was down-regulated on wheat straw. For growth on alder, no significant differential expression was observed for the putative peroxidases (Fig. 1). Of the 137 possible cytochrome P450 proteins, 33 were up- and 18 down-regulated in wheat straw, and 20 were up- and 24 down-regulated on alder (Fig. 1).

**Figure 1:**
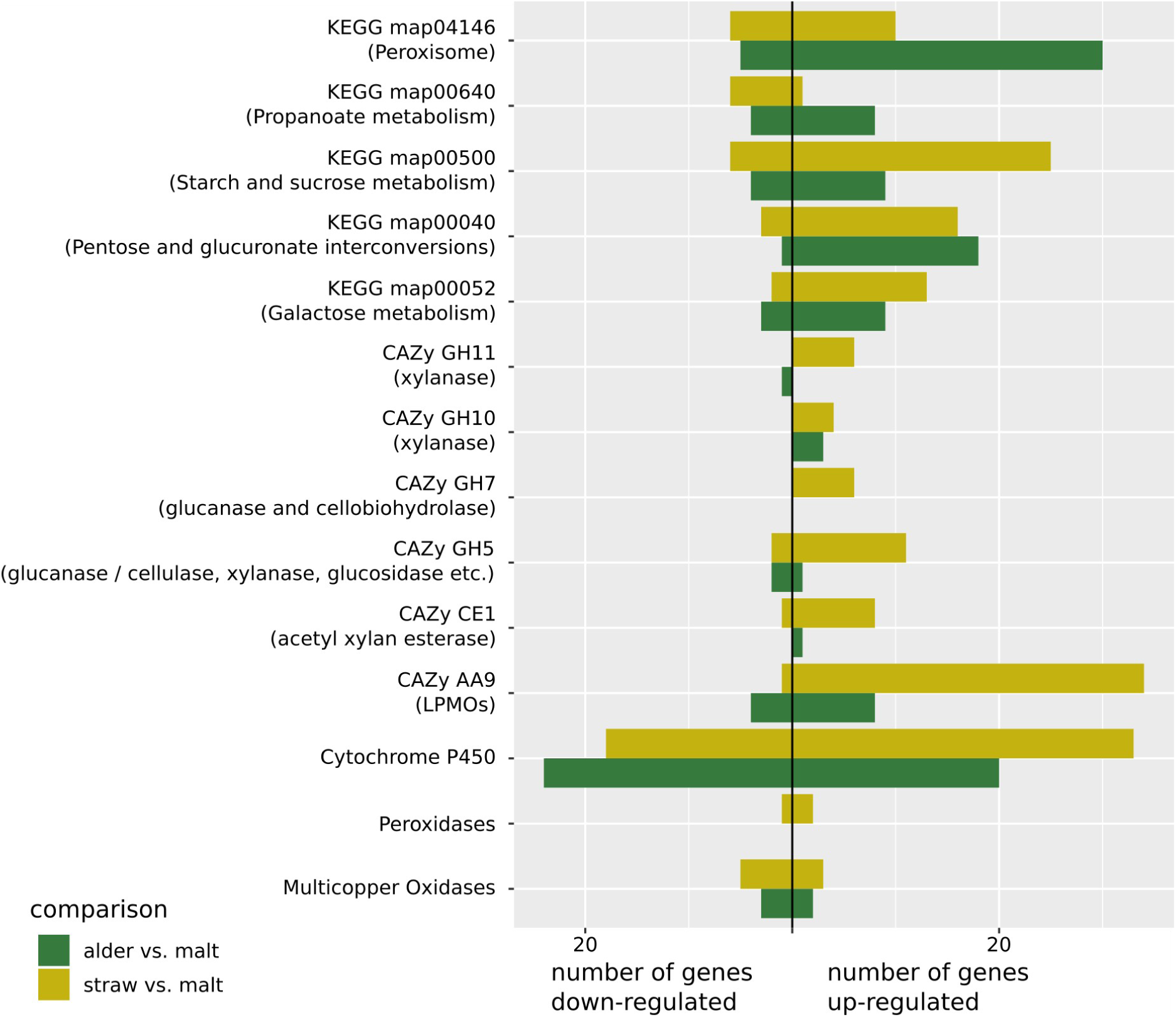
Number of up- and down-regulated genes in different gene groups for the comparison between wheat straw vs. malt extract (yellow) and between alder leaves vs. malt extract (green).

### Gene and Gene Set Activation

We found multiple activated GO terms, KEGG pathways and CAZy families for all comparisons (Supplemental Table 1-3). The only exception was the comparison between the exponential growth phase on alder leaves and on malt extract, where no active CAZy family was identified. We concentrate here on the differential expression between exponential growth on wheat straw and on malt extract, and between exponential growth on alder leaves and on malt extract, because they are the most relevant when investigating biomass degradation (Fig. 1).

From the six CAZy families that were predicted by MGSA to be regulated for growth on wheat straw (Table 3A), three (CE1, GH10 and GH11) were linked to xylane and thus hemicellulose degradation (Zhang et al., 2011) and two (GH7 and GH5_5) were linked to glucan and cellulose degradation. The CAZy family predicted to be regulated with the most genes was AA9 which contains lytic polysaccharide monooxygenases (LPMOs) acting among others on cellulose to prepare it for further enzymatic degradation and has been shown to degrade hemicellulose as well (Agger et al., 2014). Investigation of the expression of the genes assigned to these groups in the *C. aquatica* genome showed that for growth on wheat straw they were almost all strongly up-regulated, while for growth on alder in most cases (except for GH10) there was no or only weak up-regulation. For each of the families CE1 and GH7 there was one of the assigned genes, that was not up-regulated. This was also the only gene in these families predicted (by signalP) to be not secreted.

**Table 3 A:**
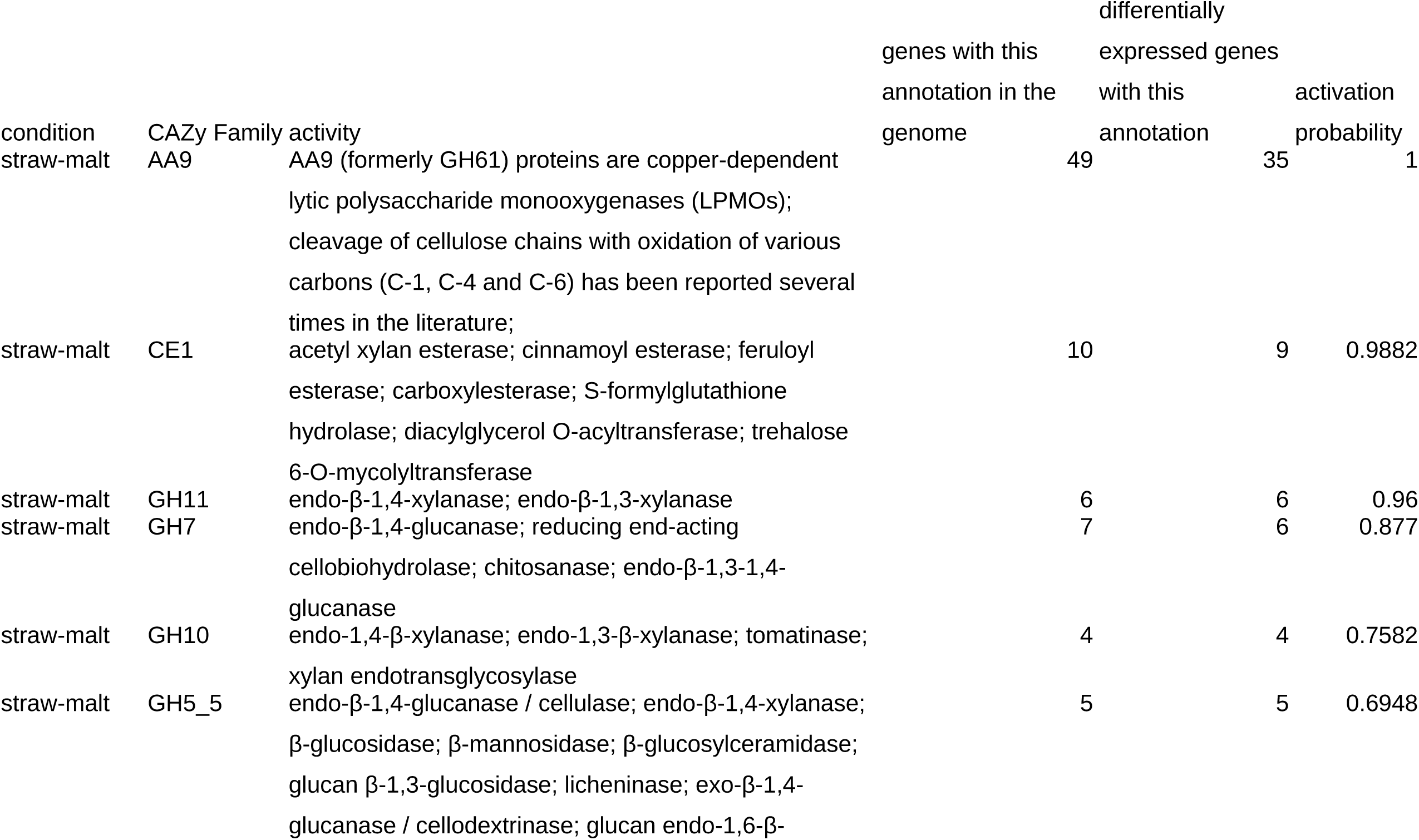
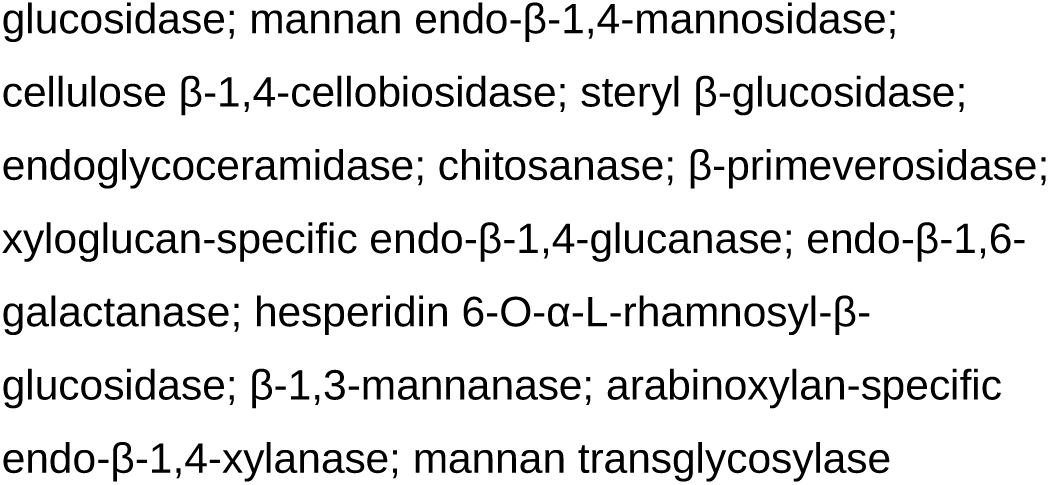
CAZy families predicted to be active by MGSA analysis. For comparison growth on straw compared to growth on malt extract

**Table 3 B:**
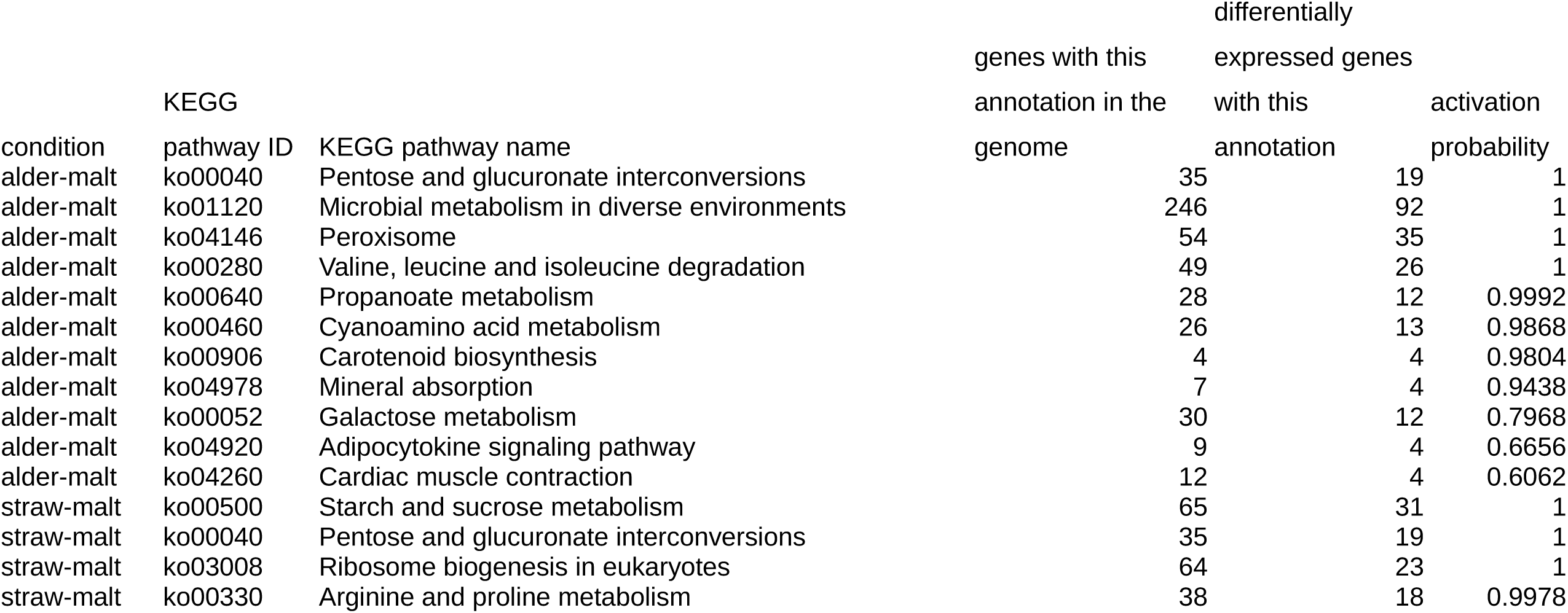
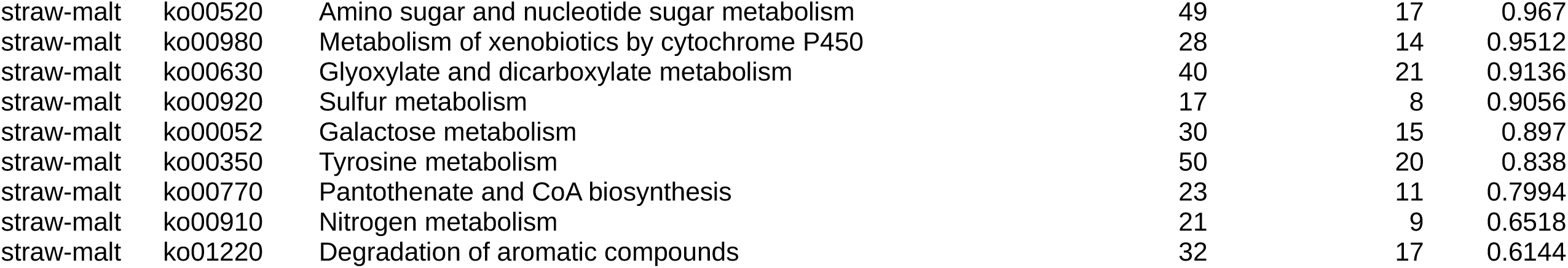
KEGG pathways predicted to be active by MGSA analysis. For comparison growth on straw compared to growth on malt extract, and growth on alder compared to malt extract

**Table 3 C:**
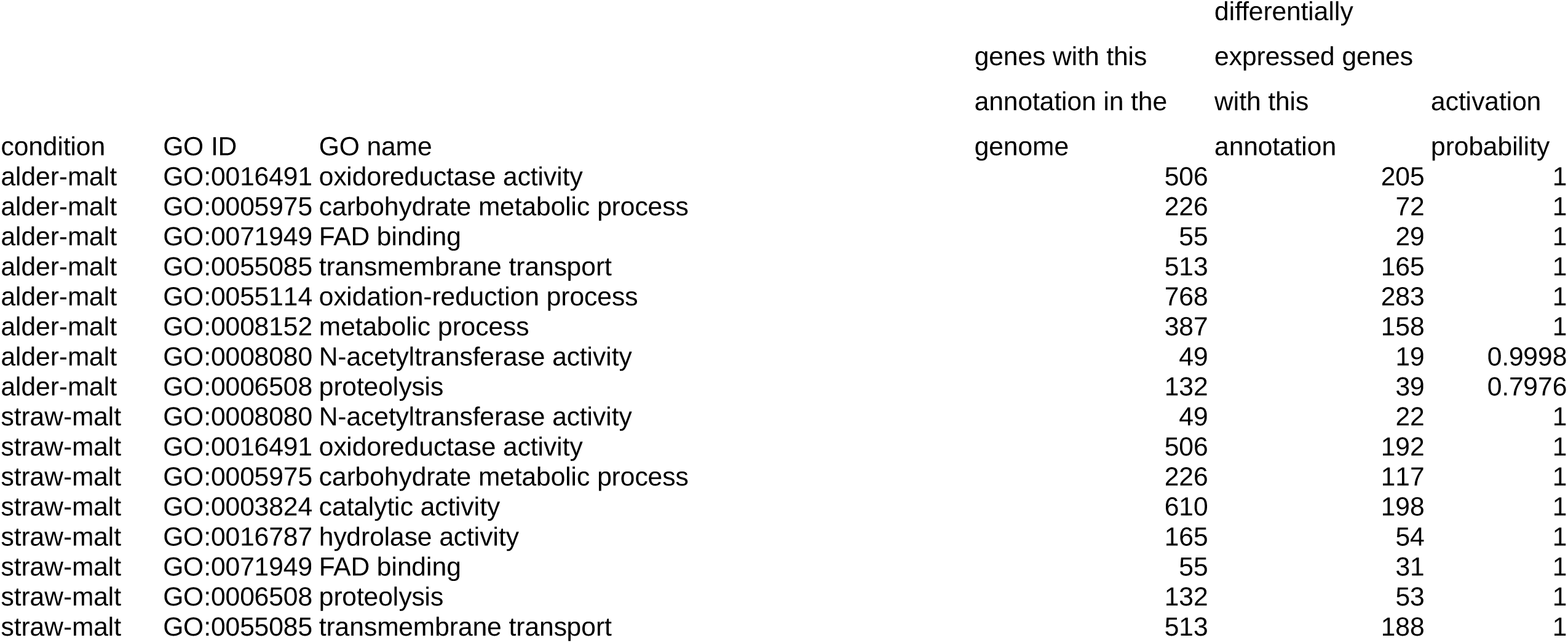
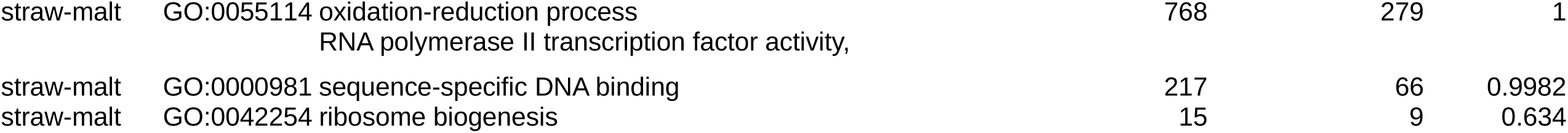
GO terms predicted to be active by MGSA analysis. For comparison growth on straw compared to growth on malt extract, and growth on alder compared to malt extract

The non-significant (Fisher’s exact test, p=0.0512) overlap between predicted activation of KEGG pathways (Table 3B) for growth on alder and wheat straw contained the two pathways ko00040 (Pentose and glucuronate interconversions) and ko00052 (Galactose metabolism). The up-regulation of the pentose and glucuronate interconversions pathway was mostly caused by the up-regulation of the genes on the path from pectin to glycerol and regulation of some genes involved in conversion of xylose to ribulose. The up-regulated enzymes in the galactose metabolism catalyze conversion of galactose into glucose. The pathway ko00500 (Starch and sucrose metabolism) was only predicted to be regulated for growth on wheat straw by the MGSA analysis. Most of the regulated genes are involved in cellulose degradation into glucose, but there is also down regulation of conversion of maltose into glucose. Although this pathway was not predicted to be regulated for growth on alder, many of the genes showed differential expression for that comparison as well. Two interesting pathways predicted to be activated for growth on alder, but not on wheat straw, were ko04146 (Peroxisome) and ko00640 (Propanoate metabolism). In ko04146, in addition to multiple genes that are important for structure and function of the peroxisome, genes involved in the β-oxidation in the peroxisome were up-regulated. In ko00640, genes for the degradation of propanoate through the β-oxidation into Acetyl-CoA were up-regulated.

The activated GO terms (Table 3C) were mostly connected to metabolism, but were not specific enough to lead to any further conclusions. GO terms predicted to be regulated in the comparison between growth on wheat straw versus growth on malt extract had a significant overlap (Fisher’s exact test, p<10^−15^) with GO terms predicted to be regulated for the comparison between growth on alder leaves versus growth on malt extract.

## Discussion

We found ten possible laccases in the genome of *C. aquatica*, of which five had previously been identified (Solé et al., 2012). The fact that one of them was assigned to the “Basidomycete Laccase” superfamily was somewhat surprising, given that *C. aquatica* is a member of the Ascomycota; however, further investigation revealed that multiple sequences from Ascomycota are already contained in this group in the database, and thus the name of the superfamily is probably just historical. The ten possible laccases exhibited both up- and down-regulation on alder and wheat straw. This result builds upon an earlier study (Solé et al., 2012) which observed differently regulated laccase genes in *C. aquatica* in response to metals and to xenobiotics and lignocellulose breakdown products, and also observed differences among growth stages. The difference we observed between alder and wheat straw may indicate that different *C. aquatica* laccases act on these substrates. The results could also be explained if different laccases are involved at different stages of *C. aquatica* growth and substrate decomposition stages, and if our cultures on wheat straw and alder were in different growth stages. This explanation seems less likely, because samples were taken at the same time and culture conditions were closely controlled.

We identified five potential peroxidases from the *C. aquatica* genome that were assigned to class II of the non-animal peroxidase superfamily by Peroxiscan. This class also contains peroxidases known to be involved in lignin degradation in Basidiomycota such as lignin peroxidase (LiP), manganese peroxidase (MnP), and versatile peroxidase (VP) (Hammel and Cullen, 2008). The peroxidases identified from the *C. aquatica* genome however, are more similar to the Ascomycota Class II family (Mathé et al., 2019). Two of these putative peroxidases were up-regulated on wheat straw, but not on alder, and to our knowledge, the expression of active peroxidase enzymes has not yet been reported for *C. aquatica*. Their activation on the presumably more lignin-rich wheat straw (compared to alder leaves; see below) could indicate that they are involved in the biotransformation of certain, perhaps phenolic lignin constituents; thus possibly contributing to the detoxification of such compounds. The enzymes of the Ascomycota Class II family do not show the specific amino acids to form the active site for lignin degradation with the same mechanisms as MnP, LiP and VP in Basidiomycota, and their function is not yet understood. Their occurrence in mostly saprotrophic fungi has lead to speculation that they might be involved in cell wall penetration and not in lignin decomposition (Mathé et al., 2019).

The third group of enzymes that we investigated belongs to the cytochrome P450 superfamily. Because the classification of these enzymes could not be further specified it is not clear which of the more than 100 enzymes from this superfamily we found. Potentially, these could act on aromatic structures stemming from lignin or other plant constituents, or on aliphatic compounds (Syed et al., 2014) such as waxes of the leaf cuticle. The observed up-regulation of some of these putative cytochrome P450 monooxygenases on wheat straw (n=33) and alder (n=20) is in favor of such functions.

We only detected clear activation of CAZy families during the exponential growth phase on wheat straw. The activated families all have cellulose- and hemicellulose-degrading activity, as expected on this substrate. The two classical glycoside hydrolase families (GH7 and GH11) that showed the highest activation probabilities have previously been reported to be induced by growth on wheat straw in other fungi (Ries et al., 2013). Besides two other glycoside hydrolase families (GH10 and GH5), which are described as acting on cellulose and hemicellulose main chain bonds, we also found up-regulation of members of the CE1 family that contains acetyl xylan esterases and of the AA9 family that contains lytic polysaccharide monooxygenases (LPMOs). Xlyan esterases cleave hemicellulose side chains, while LPMO act on cellulose and hemicellulose and have been observed to boost the conversion of lignocellulose via the oxidation of crystalline polysaccharide chains by reactive copper-superoxide complexes (Martínez et al., 2017). The AA9 is a large family and, in our case, 34 genes that were up-regulated on wheat straw have been assigned to it. Most of the targets and the specific functions of the variety of LPMOs are not yet clarified (Vaaje-Kolstad et al., 2017), but it has been shown that they cleave cellulose and hemicellulose components (Frommhagen et al., 2015). Most of the up-regulated proteins in the above mentioned CAZy families are predicted to be secreted. Together this indicates that *C. aquatica* performs extracellular degradation of cellulose and hemicellulose when grown on wheat straw. On alder leaves, none of the CAZy families were predicted as active, and very few of the genes in them were differentially expressed compared to growth on malt extract.

In contrast to this difference, the overall differentially expressed genes on wheat straw and alder showed a significant overlap and two of the KEGG pathways were regulated on both. The genes in the two common KEGG pathways (ko00040 and ko00052) encode enzymes involved in xylose and galactose degradation, and many of the genes for degradation of cellulose into glucose (ko00500) were up-regulated on both substrates as well (although many more were up-regulated on wheat straw). For growth on wheat straw, this shows a clear process of extracellular cleavage of cellulose and hemicellulose followed by utilization of the monomers as carbon sources.

We identified the up-regulation of enzymes involved in the later stages of lignocellulose degradation (e.g. xylose and galctose degradation) on both wheat straw and alder leaves, although we only detected the up-regulation of genes for the initial polymer decomposition (e.g. glycoside hydrolases, acetyl xylan esterases and LPMOs) when *C. aquatica* was grown on wheat straw. One possible explanation is the different composition of the two substrates. Wheat straw contains more cellulose (∼40%) and lignin (9-22%) (Alemdar and Sain, 2008; Bjerre et al., 1996) than alder leaves (5-15% and 6-20%, respectively) (Chauvet, 1987; Lecerf and Chauvet, 2008). It is possible that this leads to a lower expression of cell-wall degrading enzymes when *C. aquatica* is grown on alder leaves. In addition it may be that alder leaves contain carbon sources other than polysaccharides, and that these may be utilized by *C. aquatica*, which tends to grow inside the leaf matrix. Propanoate metabolism (ko00640) was predicted to be activated for growth on alder (Table 3B), and enzymes that were up-regulated in this pathway in *C. aquatica* have been found to be involved in propanoate degradation via the β-oxidation pathway in other fungi (Otzen et al., 2014). Potentially, propanoate could be produced from wax-related fatty acids in the alder leaves (for example from those of cutin and suberin not present in wheat straw), or from aliphatic side chains of plant sterols.

Gene expression of *C. aquatica* on both lignocellulose containing materials showed indication of cellulose and hemicellulose degradation. In particular the enzymes for extracellular depolymerization were more clearly up-regulated on the more cellulose rich wheat straw. Multiple laccases, peroxidases and putative cytochrome P450 monooxygenases were identified in the genome of *C. aquatica*. The expression of several of them was increased on the lignocellulose containing substrates. This observation strongly suggests that *C. aquatica* is able to modify lignin to some extent; perhaps in order to facilitate the utilization of polysaccharide components of lignocellulose as carbon and energy sources. Considering the known substrate promiscuity of the aforementioned enzymes, it further emphasizes a role of *C. aquatica* in the breakdown of xenobiotic environmental pollutants when dwelling in its natural riverine habitat.

## Acknowledgments

We would like to thank Susan Mbedi and Katharina Frindte for performing genome sequencing, Madlen Schubert for carrying out fungal cultivation, and Sajeet Hariads and Bishoy Hanna Kamel for setting up the Mycocosm portal for our assembly. We gratefully acknowledge support by the Helmholtz Association of German Research Centers under the research program “Renewable Energies”. Research was partially funded by the Leibniz Association Pakt/SAW project “MycoLink” (SAW-2014-IGB-1). RNA-Seq was funded by the Community Science Program (CSP 1663) of the Joint Genome Institute (JGI). The work conducted by the U.S. Department of Energy Joint Genome Institute, a DOE Office of Science User Facility, was supported by the Office of Science of the U.S. Department of Energy under Contract No. DE-AC02-05CH11231.

## Author Contribution

F.H., E.C.B., C.W., D.S. and M.T.M. conceived and designed the overall study. D.S. designed and oversaw cultivation. E.C.B. and E.F. planned and carried out RNA extraction, and first RNA quality checks. A.L., I.G., V.N. and G.H. performed further RNA quality assessments, RNA library preparation, and RNA sequencing. F.H. assembled and annotated the genome, performed RNA-Seq data processing and did statistical analysis. F.H., C.W., D.S. and M.T.M. wrote the intital manuscript. All authors contributed to and approved the final manuscript draft.

## References

Agger, J.W., Isaksen, T., Várnai, A., Vidal-Melgosa, S., Willats, W.G.T., Ludwig, R., Horn, S.J., Eijsink, V.G.H., Westereng, B., 2014. Discovery of LPMO activity on hemicelluloses shows the importance of oxidative processes in plant cell wall degradation. Proc Natl Acad Sci U S A 111, 6287–6292. https://doi.org/10.1073/pnas.1323629111

Alemdar, A., Sain, M., 2008. Biocomposites from wheat straw nanofibers: Morphology, thermal and mechanical properties. Composites Science and Technology 68, 557–565. https://doi.org/10.1016/j.compscitech.2007.05.044

Ashburner, M., Ball, C.A., Blake, J.A., Botstein, D., Butler, H., Cherry, J.M., Davis, A.P., Dolinski, K., Dwight, S.S., Eppig, J.T., Harris, M.A., Hill, D.P., Issel-Tarver, L., Kasarskis, A., Lewis, S., Matese, J.C., Richardson, J.E., Ringwald, M., Rubin, G.M., Sherlock, G., 2000. Gene Ontology: tool for the unification of biology. Nat Genet 25, 25–29. https://doi.org/10.1038/75556

Bauer, S., Gagneur, J., Robinson, P.N., 2010. GOing Bayesian: model-based gene set analysis of genome-scale data. Nucleic Acids Res 38, 3523–3532. https://doi.org/10.1093/nar/gkq045

Bauer, S., Robinson, P.N., Gagneur, J., 2011. Model-based gene set analysis for Bioconductor. Bioinformatics 27, 1882–1883. https://doi.org/10.1093/bioinformatics/btr296

Bjerre, A.B., Olesen, A.B., Fernqvist, T., Plöger, A., Schmidt, A.S., 1996. Pretreatment of wheat straw using combined wet oxidation and alkaline hydrolysis resulting in convertible cellulose and hemicellulose. Biotechnology and Bioengineering 49, 568–577. https://doi.org/10.1002/(SICI)1097-0290(19960305)49:5<568::AID-BIT10>3.0.CO;2-6

Bourne, E.C., Johnston, P.R., Funke, E., Monaghan, M.T., in press. Gene expression analysis of litter-associated fungi using RNA-seq, in: Methods to Study Litter Decomposition. Springer.

Brown, C.T., Howe, A., Zhang, Q., Pyrkosz, A.B., Brom, T.H., 2012. A reference-free algorithm for computational normalization of shotgun sequencing data. 1203.4802 [q-bio].

Chauvet, E., 1987. Changes in the chemical composition of alder, poplar and willow leaves during decomposition in a river. Hydrobiologia 148, 35–44. https://doi.org/10.1007/BF00018164

Crusoe, M.R., Alameldin, H.F., Awad, S., Boucher, E., Caldwell, A., Cartwright, R., Charbonneau, A., Constantinides, B., Edvenson, G., Fay, S., Fenton, J., Fenzl, T., Fish, J., Garcia-Gutierrez, L., Garland, P., Gluck, J., González, I., Guermond, S., Guo, J., Gupta, A., Herr, J.R., Howe, A., Hyer, A., Härpfer, A., Irber, L., Kidd, R., Lin, D., Lippi, J., Mansour, T., McA’Nulty, P., McDonald, E., Mizzi, J., Murray, K.D., Nahum, J.R., Nanlohy, K., Nederbragt, A.J., Ortiz-Zuazaga, H., Ory, J., Pell, J., Pepe-Ranney, C., Russ, Z.N., Schwarz, E., Scott, C., Seaman, J., Sievert, S., Simpson, J., Skennerton, C.T., Spencer, J., Srinivasan, R., Standage, D., Stapleton, J.A., Steinman, S.R., Stein, J., Taylor, B., Trimble, W., Wiencko, H.L., Wright, M., Wyss, B., Zhang, Q., zyme, en, Brown, C.T., 2015. The khmer software package: enabling efficient nucleotide sequence analysis. F1000Res 4. https://doi.org/10.12688/f1000research.6924.1

Delmas, S., Pullan, S.T., Gaddipati, S., Kokolski, M., Malla, S., Blythe, M.J., Ibbett, R., Campbell, M., Liddell, S., Aboobaker, A., Tucker, G.A., Archer, D.B., 2012. Uncovering the genome-wide transcriptional responses of the filamentous fungus Aspergillus niger to lignocellulose using RNA sequencing. PLOS Genetics 8, e1002875. https://doi.org/10.1371/journal.pgen.1002875

Dobin, A., Davis, C.A., Schlesinger, F., Drenkow, J., Zaleski, C., Jha, S., Batut, P., Chaisson, M., Gingeras, T.R., 2013. STAR: ultrafast universal RNA-seq aligner. Bioinformatics 29, 15–21. https://doi.org/10.1093/bioinformatics/bts635

Duarte, S., Bärlocher, F., Trabulo, J., Cássio, F., Pascoal, C., 2015. Stream-dwelling fungal decomposer communities along a gradient of eutrophication unraveled by 454 pyrosequencing. Fungal Diversity 70, 127–148. https://doi.org/10.1007/s13225-014-0300-y

Fawal, N., Li, Q., Savelli, B., Brette, M., Passaia, G., Fabre, M., Mathé, C., Dunand, C., 2013. PeroxiBase: a database for large-scale evolutionary analysis of peroxidases. Nucleic Acids Res 41, D441–D444. https://doi.org/10.1093/nar/gks1083

Finn, R.D., Attwood, T.K., Babbitt, P.C., Bateman, A., Bork, P., Bridge, A.J., Chang, H.-Y., Dosztányi, Z., El-Gebali, S., Fraser, M., Gough, J., Haft, D., Holliday, G.L., Huang, H., Huang, X., Letunic, I., Lopez, R., Lu, S., Marchler-Bauer, A., Mi, H., Mistry, J., Natale, D.A., Necci, M., Nuka, G., Orengo, C.A., Park, Y., Pesseat, S., Piovesan, D., Potter, S.C., Rawlings, N.D., Redaschi, N., Richardson, L., Rivoire, C., Sangrador-Vegas, A., Sigrist, C., Sillitoe, I., Smithers, B., Squizzato, S., Sutton, G., Thanki, N., Thomas, P.D., Tosatto, S.C.E., Wu, C.H., Xenarios, I., Yeh, L.-S., Young, S.-Y., Mitchell, A.L., 2017. InterPro in 2017-beyond protein family and domain annotations. Nucleic Acids Res. 45, D190–D199. https://doi.org/10.1093/nar/gkw1107

Floudas, D., Binder, M., Riley, R., Barry, K., Blanchette, R.A., Henrissat, B., Martínez, A.T., Otillar, R., Spatafora, J.W., Yadav, J.S., Aerts, A., Benoit, I., Boyd, A., Carlson, A., Copeland, A., Coutinho, P.M., Vries, R.P. de, Ferreira, P., Findley, K., Foster, B., Gaskell, J., Glotzer, D., Górecki, P., Heitman, J., Hesse, C., Hori, C., Igarashi, K., Jurgens, J.A., Kallen, N., Kersten, P., Kohler, A., Kües, U., Kumar, T.K.A., Kuo, A., LaButti, K., Larrondo, L.F., Lindquist, E., Ling, A., Lombard, V., Lucas, S., Lundell, T., Martin, R., McLaughlin, D.J., Morgenstern, I., Morin, E., Murat, C., Nagy, L.G., Nolan, M., Ohm, R.A., Patyshakuliyeva, A., Rokas, A., Ruiz-Dueñas, F.J., Sabat, G., Salamov, A., Samejima, M., Schmutz, J., Slot, J.C., John, F.S., Stenlid, J., Sun, H., Sun, S., Syed, K., Tsang, A., Wiebenga, A., Young, D., Pisabarro, A., Eastwood, D.C., Martin, F., Cullen, D., Grigoriev, I.V., Hibbett, D.S., 2012. The paleozoic origin of enzymatic lignin decomposition reconstructed from 31 fungal genomes. Science 336, 1715–1719. https://doi.org/10.1126/science.1221748

Frommhagen, M., Mutte, S.K., Westphal, A.H., Koetsier, M.J., Hinz, S.W.A., Visser, J., Vincken, J.-P., Weijers, D., van Berkel, W.J.H., Gruppen, H., Kabel, M.A., 2017. Boosting LPMO-driven lignocellulose degradation by polyphenol oxidase-activated lignin building blocks. Biotechnology for Biofuels 10, 121. https://doi.org/10.1186/s13068-017-0810-4

Frommhagen, M., Sforza, S., Westphal, A.H., Visser, J., Hinz, S.W.A., Koetsier, M.J., van Berkel, W.J.H., Gruppen, H., Kabel, M.A., 2015. Discovery of the combined oxidative cleavage of plant xylan and cellulose by a new fungal polysaccharide monooxygenase. Biotechnology for Biofuels 8, 101. https://doi.org/10.1186/s13068-015-0284-1

Gessner, M.O., Gulis, K.A., E. Kuehn, Chauvet, E., Suberkopp, K., 2007. Fungal decomposers of plant litter in aquatic ecosystems., in: Environmental and Microbial Relationships, The Mycota. Springer, Berlin, Heidelberg, pp. 301–324.

Grabherr, M.G., Haas, B.J., Yassour, M., Levin, J.Z., Thompson, D.A., Amit, I., Adiconis, X., Fan, L., Raychowdhury, R., Zeng, Q., Chen, Z., Mauceli, E., Hacohen, N., Gnirke, A., Rhind, N., di Palma, F., Birren, B.W., Nusbaum, C., Lindblad-Toh, K., Friedman, N., Regev, A., 2011. Trinity: reconstructing a full-length transcriptome without a genome from RNA-Seq data. Nat Biotechnol 29, 644–652. https://doi.org/10.1038/nbt.1883

Grigoriev, I.V., Nikitin, R., Haridas, S., Kuo, A., Ohm, R., Otillar, R., Riley, R., Salamov, A., Zhao, X., Korzeniewski, F., Smirnova, T., Nordberg, H., Dubchak, I., Shabalov, I., 2014. MycoCosm portal: gearing up for 1000 fungal genomes. Nucleic Acids Res. 42, D699–704. https://doi.org/10.1093/nar/gkt1183

Haas, B.J., Delcher, A.L., Mount, S.M., Wortman, J.R., Smith, R.K., Hannick, L.I., Maiti, R., Ronning, C.M., Rusch, D.B., Town, C.D., Salzberg, S.L., White, O., 2003. Improving the Arabidopsis genome annotation using maximal transcript alignment assemblies. Nucleic Acids Res 31, 5654–5666. https://doi.org/10.1093/nar/gkg770

Hammel, K.E., Cullen, D., 2008. Role of fungal peroxidases in biological ligninolysis. Current Opinion in Plant Biology, Physiology and Metabolism - Edited by Markus Pauly and Kenneth Keegstra 11, 349–355. https://doi.org/10.1016/j.pbi.2008.02.003

Iqbal, S.H., Webster, J., 1973. Aquatic hyphomycete spora of the River Exe and its tributaries. Transactions of the British Mycological Society 61, 331–346. https://doi.org/10.1016/S0007-1536(73)80155-X

Johnson, M.T.J., Carpenter, E.J., Tian, Z., Bruskiewich, R., Burris, J.N., Carrigan, C.T., Chase, M.W., Clarke, N.D., Covshoff, S., dePamphilis, C.W., Edger, P.P., Goh, F., Graham, S., Greiner, S., Hibberd, J.M., Jordon-Thaden, I., Kutchan, T.M., Leebens-Mack, J., Melkonian, M., Miles, N., Myburg, H., Patterson, J., Pires, J.C., Ralph, P., Rolf, M., Sage, R.F., Soltis, D., Soltis, P., Stevenson, D., Jr, C.N.S. Surek, B., Thomsen, C.J.M., Villarreal, J.C., Wu, X., Zhang, Y., Deyholos, M.K., Wong, G.K.-S., 2012. Evaluating methods for isolating total RNA and predicting the success of sequencing phylogenetically diverse plant transcriptomes. PLOS ONE 7, e50226. https://doi.org/10.1371/journal.pone.0050226

Jon Palmer, Jason Stajich, 2019. nextgenusfs/funannotate: funannotate v1.5.3. Zenodo. https://doi.org/10.5281/zenodo.2604804

Jones, P., Binns, D., Chang, H.-Y., Fraser, M., Li, W., McAnulla, C., McWilliam, H., Maslen, J., Mitchell, A., Nuka, G., Pesseat, S., Quinn, A.F., Sangrador-Vegas, A., Scheremetjew, M., Yong, S.-Y., Lopez, R., Hunter, S., 2014. InterProScan 5: genome-scale protein function classification. Bioinformatics 30, 1236–1240. https://doi.org/10.1093/bioinformatics/btu031

Junghanns, C., Moeder, M., Krauss, G., Martin, C., Schlosser, D., 2005. Degradation of the xenoestrogen nonylphenol by aquatic fungi and their laccases. Microbiology (Reading, Engl.) 151, 45–57. https://doi.org/10.1099/mic.0.27431-0

Kanehisa, M., Sato, Y., Kawashima, M., Furumichi, M., Tanabe, M., 2016a. KEGG as a reference resource for gene and protein annotation. Nucleic Acids Res. 44, D457–462. https://doi.org/10.1093/nar/gkv1070

Kanehisa, M., Sato, Y., Morishima, K., 2016b. BlastKOALA and GhostKOALA: KEGG tools for functional characterization of genome and metagenome sequences. J. Mol. Biol. 428, 726–731. https://doi.org/10.1016/j.jmb.2015.11.006

Köster, J., Rahmann, S., 2012. Snakemake—a scalable bioinformatics workflow engine. Bioinformatics 28, 2520–2522. https://doi.org/10.1093/bioinformatics/bts480

Koua, D., Cerutti, L., Falquet, L., Sigrist, C.J.A., Theiler, G., Hulo, N., Dunand, C., 2009. PeroxiBase: a database with new tools for peroxidase family classification. Nucleic Acids Res. 37, D261–266. https://doi.org/10.1093/nar/gkn680

Krauss, G.-J., Solé, M., Krauss, G., Schlosser, D., Wesenberg, D., Bärlocher, F., 2011. Fungi in freshwaters: ecology, physiology and biochemical potential. FEMS Microbiol. Rev. 35, 620–651. https://doi.org/10.1111/j.1574-6976.2011.00266.x

Kumar, S.V.S., Phale, P.S., Durani, S., Wangikar, P.P., 2003. Combined sequence and structure analysis of the fungal laccase family. Biotechnol. Bioeng. 83, 386–394. https://doi.org/10.1002/bit.10681

Lecerf, A., Chauvet, E., 2008. Intraspecific variability in leaf traits strongly affects alder leaf decomposition in a stream. Basic and Applied Ecology 9, 598–605. https://doi.org/10.1016/j.baae.2007.11.003

Li, B., Dewey, C.N., 2011. RSEM: accurate transcript quantification from RNA-Seq data with or without a reference genome. BMC Bioinformatics 12, 323. https://doi.org/10.1186/1471-2105-12-323

Lombard, V., Golaconda Ramulu, H., Drula, E., Coutinho, P.M., Henrissat, B., 2014. The carbohydrate-active enzymes database (CAZy) in 2013. Nucleic Acids Res. 42, D490–495. https://doi.org/10.1093/nar/gkt1178

Love, M.I., Huber, W., Anders, S., 2014. Moderated estimation of fold change and dispersion for RNA-seq data with DESeq2. Genome Biol. 15, 550. https://doi.org/10.1186/s13059-014-0550-8

Lundell, T.K., Mäkelä, M.R., Hildén, K., 2010. Lignin-modifying enzymes in filamentous basidiomycetes--ecological, functional and phylogenetic review. J. Basic Microbiol. 50, 5–20. https://doi.org/10.1002/jobm.200900338

Martin, C., Moeder, M., Daniel, X., Krauss, G., Schlosser, D., 2007. Biotransformation of the polycyclic musks HHCB and AHTN and metabolite formation by fungi occurring in freshwater environments. Environ. Sci. Technol. 41, 5395–5402. https://doi.org/10.1021/es0711462

Martínez, A.T., Ruiz-Dueñas, F.J., Camarero, S., Serrano, A., Linde, D., Lund, H., Vind, J., Tovborg, M., Herold-Majumdar, O.M., Hofrichter, M., Liers, C., Ullrich, R., Scheibner, K., Sannia, G., Piscitelli, A., Pezzella, C., Sener, M.E., Kılıç, S., van Berkel, W.J.H., Guallar, V., Lucas, M.F., Zuhse, R., Ludwig, R., Hollmann, F., Fernández-Fueyo, E., Record, E., Faulds, C.B., Tortajada, M., Winckelmann, I., Rasmussen, J.-A., Gelo-Pujic, M., Gutiérrez, A., del Río, J.C., Rencoret, J., Alcalde, M., 2017. Oxidoreductases on their way to industrial biotransformations. Biotechnology Advances 35, 815–831. https://doi.org/10.1016/j.biotechadv.2017.06.003

Mathé, C., Fawal, N., Roux, C., Dunand, C., 2019. In silico definition of new ligninolytic peroxidase sub-classes in fungi and putative relation to fungal life style. Sci Rep 9, 1–14. https://doi.org/10.1038/s41598-019-56774-4

Otzen, C., Bardl, B., Jacobsen, I.D., Nett, M., Brock, M., 2014. Candida albicans utilizes a modified β-oxidation pathway for the degradation of toxic propionyl-CoA. J. Biol. Chem. 289, 8151–8169. https://doi.org/10.1074/jbc.M113.517672

Petersen, T.N., Brunak, S., Heijne, G. von, Nielsen, H., 2011. SignalP 4.0: discriminating signal peptides from transmembrane regions. Nat Methods 8, 785–786. https://doi.org/10.1038/nmeth.1701

Ries, L., Pullan, S.T., Delmas, S., Malla, S., Blythe, M.J., Archer, D.B., 2013. Genome-wide transcriptional response of Trichoderma reesei to lignocellulose using RNA sequencing and comparison with Aspergillus niger. BMC Genomics 14, 541. https://doi.org/10.1186/1471-2164-14-541

Riley, R., Salamov, A.A., Brown, D.W., Nagy, L.G., Floudas, D., Held, B.W., Levasseur, A., Lombard, V., Morin, E., Otillar, R., Lindquist, E.A., Sun, H., LaButti, K.M., Schmutz, J., Jabbour, D., Luo, H., Baker, S.E., Pisabarro, A.G., Walton, J.D., Blanchette, R.A., Henrissat, B., Martin, F., Cullen, D., Hibbett, D.S., Grigoriev, I.V., 2014. Extensive sampling of basidiomycete genomes demonstrates inadequacy of the white-rot/brown-rot paradigm for wood decay fungi. PNAS 111, 9923–9928. https://doi.org/10.1073/pnas.1400592111

Schlosser, D., Höfer, C., 2002. Laccase-catalyzed oxidation of Mn2+ in the presence of natural Mn3+ chelators as a novel source of extracellular H2O2 production and its impact on manganese peroxidase. Appl. Environ. Microbiol. 68, 3514–3521. https://doi.org/10.1128/AEM.68.7.3514-3521.2002

Simão, F.A., Waterhouse, R.M., Ioannidis, P., Kriventseva, E.V., Zdobnov, E.M., 2015. BUSCO: assessing genome assembly and annotation completeness with single-copy orthologs. Bioinformatics 31, 3210–3212. https://doi.org/10.1093/bioinformatics/btv351

Sirim, D., Wagner, F., Wang, L., Schmid, R.D., Pleiss, J., 2011. The Laccase Engineering Database: a classification and analysis system for laccases and related multicopper oxidases. Database (Oxford) 2011. https://doi.org/10.1093/database/bar006

Solé, M., Müller, I., Pecyna, M.J., Fetzer, I., Harms, H., Schlosser, D., 2012. Differential regulation by organic compounds and heavy metals of multiple laccase genes in the aquatic hyphomycete Clavariopsis aquatica. Appl. Environ. Microbiol. 78, 4732–4739. https://doi.org/10.1128/AEM.00635-12

Suberkropp, K., Klug, M.J., 1976. Fungi and bacteria associated with leaves during processing in a woodland stream. Ecology 57, 707–719. https://doi.org/10.2307/1936184

Syed, K., Shale, K., Pagadala, N.S., Tuszynski, J., 2014. Systematic Identification and Evolutionary Analysis of Catalytically Versatile Cytochrome P450 Monooxygenase Families Enriched in Model Basidiomycete Fungi. PLOS ONE 9, e86683. https://doi.org/10.1371/journal.pone.0086683

Tang, J.D., Parker, L.A., Perkins, A.D., Sonstegard, T.S., Schroeder, S.G., Nicholas, D.D., Diehl, S.V., 2013. Gene expression analysis of copper tolerance and wood decay in the brown rot fungus Fibroporia radiculosa. Appl. Environ. Microbiol. 79, 1523–1533. https://doi.org/10.1128/AEM.02916-12

The Gene Ontology Consortium, 2017. Expansion of the Gene Ontology knowledgebase and resources. Nucleic Acids Res 45, D331–D338. https://doi.org/10.1093/nar/gkw1108

Vaaje-Kolstad, G., Forsberg, Z., Loose, J.S., Bissaro, B., Eijsink, V.G., 2017. Structural diversity of lytic polysaccharide monooxygenases. Current Opinion in Structural Biology, Carbohydrates: A feast of structural glycobiology • Sequences and topology: Computational studies of protein-protein interactions 44, 67–76. https://doi.org/10.1016/j.sbi.2016.12.012

Yang, Y., Fan, F., Zhuo, R., Ma, F., Gong, Y., Wan, X., Jiang, M., Zhang, X., 2012. Expression of the laccase gene from a white rot fungus in Pichia pastoris can enhance the resistance of this yeast to H2O2-mediated oxidative stress by stimulating the glutathione-based antioxidative system. Appl. Environ. Microbiol. 78, 5845–5854. https://doi.org/10.1128/AEM.00218-12

Yin, Y., Mao, X., Yang, J., Chen, X., Mao, F., Xu, Y., 2012. dbCAN: a web resource for automated carbohydrate-active enzyme annotation. Nucleic Acids Res. 40, W445–451. https://doi.org/10.1093/nar/gks479

Zerbino, D.R., Birney, E., 2008. Velvet: algorithms for de novo short read assembly using de Bruijn graphs. Genome Res. 18, 821–829. https://doi.org/10.1101/gr.074492.107

Zhang, J., Siika-Aho, M., Tenkanen, M., Viikari, L., 2011. The role of acetyl xylan esterase in the solubilization of xylan and enzymatic hydrolysis of wheat straw and giant reed. Biotechnol Biofuels 4, 60. https://doi.org/10.1186/1754-6834-4-60

Zhang, Q., Pell, J., Canino-Koning, R., Howe, A.C., Brown, C.T., 2014. These Are Not the K-mers You Are Looking For: Efficient Online K-mer Counting Using a Probabilistic Data Structure. PLOS ONE 9, e101271. https://doi.org/10.1371/journal.pone.0101271

