## Supplementary material for "Identification of enzymes for lignocellulose degradation in the genome and transcriptome of the aquatic hyphomycete *Clavariopsis aquatica*": Supllemental Info

### Supplemental Info 1: Commands

**Digital normalization** was done with the khmer package with the following steps and parameters:

1. `interleave-reads.py`
2. `normalize-by-median.py -C 20 -k 20 -N 4 -x 2.5e8`
3. `filter-abund.py clavariopsis.kh`
4. `normalize-by-median.py -C 5 -k 20 -N 4 -x 1e8`
5. `extract-paired-reads.py`

**Genome assembly** was done was done with velvet with the digitally normalized reads with the following commands:

1. `velveth -fastq.gz -shortPaired`
2. `velvetg -ins_length 900 -exp_cov 140 -cov_cutoff 50`

***De Novo* transcriptome assembly** was done with Trinity with the following command:

```
Trinity --seqType fq --max_memory 100G --single $INDATA --CPU 12
--trimmomatic --normalize_reads --SS_lib_type R
```

**Genome guided transcriptome assembly** was done with Trinity with the following command:

```
Trinity --genome_guided_bam $INDATA --genome_guided_max_intron
1000 --max_memory 30G --CPU 12
```

The **PASA pipeline** was run with the following commands:

1. `cat Trinity.fasta Trinity-GG.fasta > ${TRANSCRIPTS}`
2. `accession_extractor.pl < Trinity.fasta > tdn.accs`

3. `Launch_PASA_pipeline.pl -c alignAssembly.conf -C -R -g ${GENOME} -t ${TRANSCRIPTS} --ALIGNERS blat,gmap --TDN tdn.accs --transcribed_is_aligned_orient --MAX_INTRON_LENGTH 3000`
4. `build_comprehensive_transcriptome.dbi -c alignAssembly.conf -t ${TRANSCRIPTS} --min_per_ID 95 --min_per_aligned 30`

### Supplemental Table 1

| condition | CAZy Family | activity | genes with this annotation in the genome | differentially expressed genes with this annotation | activation probability |
|---|---|---|---|---|---|
| alder-straw | AA9 | AA9 (formerly GH61) proteins are copper-dependent lytic polysaccharide monooxygenases (LPMOs); cleavage of cellulose chains with oxidation of various carbons (C-1, C-4 and C-6) has been reported several times in the literature; | 49 | 35 | 1 |
| alder-straw | CE1 | acetyl xylan esterase; cinnamoyl esterase; feruloyl esterase; carboxylesterase; S-formylglutathione hydrolase; diacylglycerol O-acyltransferase; trehalose 6-O-mycolyltransferase | 10 | 9 | 0.9826 |
| alder-straw | GH11 | endo-β-1,4-xylanase; endo-β-1,3-xylanase | 6 | 6 | 0.935 |
| alder-straw | GH7 | endo-β-1,4-glucanase; reducing end-acting cellobiohydrolase; chitosanase; endo-β-1,3-1,4-glucanase | 7 | 6 | 0.8518 |
| alder-straw | GH131 | broad specificity exo-β-1,3/1,6-glucanase with endo-β-1,4-glucanase activity; | 4 | 4 | 0.7446 |
| alder-straw | GH10 | endo-1,4-β-xylanase; endo-1,3-β-xylanase; tomatinase; xylan endotransglycosylase | 4 | 4 | 0.7428 |
| alder-straw | GH5_5 | endo-β-1,4-glucanase / cellulase; endo-β-1,4-xylanase; β-glucosidase; β-mannosidase; β-glucosylceramidase; glucan β-1,3-glucosidase; licheninase; exo-β-1,4-glucanase / cellodextrinase; glucan endo-1,6-β-glucosidase; mannan endo-β-1,4-mannosidase; cellulose β-1,4-cellobiosidase; steryl β-glucosidase; endoglycoceramidase; chitosanase; β-primeverosidase; xyloglucan-specific endo-β-1,4-glucanase; endo-β-1,6-galactanase; hesperidin 6-O-α-L-rhamnosyl-β-glucosidase; β-1,3-mannanase; arabinoxylan- | 5 | 5 | 0.6984 |

| | | | | | |
|---|---|---|---|---|---|
| | | specific endo-β-1,4-xylanase; mannan transglycosylase | | | |
| solid-liquidExp | GH7 | endo-β-1,4-glucanase; reducing end-acting cellobiohydrolase; chitosanase; endo-β-1,3-1,4-glucanase | 7 | 7 | 0.937 |
| solid-liquidExp | AA9 | AA9 (formerly GH61) proteins are copper-dependent lytic polysaccharide monooxygenases (LPMOs); cleavage of cellulose chains with oxidation of various carbons (C-1, C-4 and C-6) has been reported several times in the literature; | 49 | 26 | 0.813 |
| solid-liquidExp | GH72 | β-1,3-glucanosyltransglycosylase | 7 | 6 | 0.7628 |
| solid-liquidExp | AA7 | glucooligosaccharide oxidase; chitooligosaccharide oxidase | 25 | 14 | 0.6684 |
| solid-liquidExp | GH3 | β-glucosidase; xylan 1,4-β-xylosidase; β-glucosylceramidase; β-N-acetylhexosaminidase; α-L-arabinofuranosidase; glucan 1,3-β-glucosidase; glucan 1,4-β-glucosidase; isoprimeverose-producing oligoxyloglucan hydrolase; coniferin β-glucosidase; exo-1,3-1,4-glucanase; β-N-acetylglucosaminide phosphorylases | 19 | 11 | 0.6038 |
| solid-liquidSta | AA9 | AA9 (formerly GH61) proteins are copper-dependent lytic polysaccharide monooxygenases (LPMOs); cleavage of cellulose chains with oxidation of various carbons (C-1, C-4 and C-6) has been reported several times in the literature; | 49 | 33 | 1 |
| solid-liquidSta | GH7 | endo-β-1,4-glucanase; reducing end-acting cellobiohydrolase; chitosanase; endo-β-1,3-1,4-glucanase | 7 | 7 | 0.9634 |
| solid-liquidSta | CE16 | acetylesterase active on various carbohydrate acetyl esters | 5 | 5 | 0.847 |
| solid-liquidSta | AA12 | The pyrroloquinoline quinone-dependent | 6 | 5 | 0.745 |

| | | | | | |
|---|---|---|---|---|---|
| | | oxidoreductase activity was demonstrated for the CC1G_09525 protein of Coprinopsis cinerea. | | | |
| solid-liquidSta | GH55 | exo-β-1,3-glucanase; endo-β-1,3-glucanase | 4 | 4 | 0.7364 |
| solid-liquidSta | GH11 | endo-β-1,4-xylanase; endo-β-1,3-xylanase | 6 | 5 | 0.7024 |
| stat-exp | CE8 | pectin methylesterase | 6 | 5 | 0.6276 |
| straw-malt | AA9 | AA9 (formerly GH61) proteins are copper-dependent lytic polysaccharide monooxygenases (LPMOs); cleavage of cellulose chains with oxidation of various carbons (C-1, C-4 and C-6) has been reported several times in the literature; | 49 | 35 | 1 |
| straw-malt | CE1 | acetyl xylan esterase; cinnamoyl esterase; feruloyl esterase; carboxylesterase; S-formylglutathione hydrolase; diacylglycerol O-acyltransferase; trehalose 6-O-mycolyltransferase | 10 | 9 | 0.9882 |
| straw-malt | GH11 | endo-β-1,4-xylanase; endo-β-1,3-xylanase | 6 | 6 | 0.96 |
| straw-malt | GH7 | endo-β-1,4-glucanase; reducing end-acting cellobiohydrolase; chitosanase; endo-β-1,3-1,4-glucanase | 7 | 6 | 0.877 |
| straw-malt | GH10 | endo-1,4-β-xylanase; endo-1,3-β-xylanase; tomatinase; xylan endotransglycosylase | 4 | 4 | 0.7582 |
| straw-malt | GH5_5 | endo-β-1,4-glucanase / cellulase; endo-β-1,4-xylanase; β-glucosidase; β-mannosidase; β-glucosylceramidase; glucan β-1,3-glucosidase; licheninase; exo-β-1,4-glucanase / cellodextrinase; glucan endo-1,6-β-glucosidase; mannan endo-β-1,4-mannosidase; cellulose β-1,4-cellobiosidase; steryl β-glucosidase; endoglycoceramidase; chitosanase; β-primeverosidase; xyloglucan-specific endo-β-1,4-glucanase; endo-β-1,6-galactanase; hesperidin 6-O-α-L-rhamnosyl-β-glucosidase; β-1,3-mannanase; arabinoxylan- | 5 | 5 | 0.6948 |

| | | |
|---|---|---|
| | | specific endo-β-1,4-xylanase; mannan transglycosylase |

### Supplemental Table 2

| condition | KEGG pathway ID | KEGG pathway name | genes with this annotation in the genome | differentially expressed genes with this annotation | activation probability |
|---|---|---|---|---|---|
| alder-malt | ko00040 | Pentose and glucuronate interconversions | 35 | 19 | 1 |
| alder-malt | ko01120 | Microbial metabolism in diverse environments | 246 | 92 | 1 |
| alder-malt | ko04146 | Peroxisome | 54 | 35 | 1 |
| alder-malt | ko00280 | Valine, leucine and isoleucine degradation | 49 | 26 | 1 |
| alder-malt | ko00640 | Propanoate metabolism | 28 | 12 | 0.9992 |
| alder-malt | ko00460 | Cyanoamino acid metabolism | 26 | 13 | 0.9868 |
| alder-malt | ko00906 | Carotenoid biosynthesis | 4 | 4 | 0.9804 |
| alder-malt | ko04978 | Mineral absorption | 7 | 4 | 0.9438 |
| alder-malt | ko00052 | Galactose metabolism | 30 | 12 | 0.7968 |
| alder-malt | ko04920 | Adipocytokine signaling pathway | 9 | 4 | 0.6656 |
| alder-malt | ko04260 | Cardiac muscle contraction | 12 | 4 | 0.6062 |
| alder-straw | ko00500 | Starch and sucrose metabolism | 65 | 32 | 1 |
| alder-straw | ko01120 | Microbial metabolism in diverse environments | 246 | 96 | 1 |
| alder-straw | ko04146 | Peroxisome | 54 | 33 | 1 |
| alder-straw | ko00040 | Pentose and glucuronate interconversions | 35 | 16 | 0.9972 |

| | | | | | |
|---|---|---|---|---|---|
| alder-straw | ko00190 | Oxidative phosphorylation | 73 | 25 | 0.9796 |
| alder-straw | ko00520 | Amino sugar and nucleotide sugar metabolism | 49 | 18 | 0.974 |
| alder-straw | ko00770 | Pantothenate and CoA biosynthesis | 23 | 10 | 0.9574 |
| alder-straw | ko04142 | Lysosome | 40 | 13 | 0.955 |
| alder-straw | ko00330 | Arginine and proline metabolism | 38 | 17 | 0.9466 |
| alder-straw | ko00906 | Carotenoid biosynthesis | 4 | 4 | 0.9202 |
| alder-straw | ko00600 | Sphingolipid metabolism | 21 | 11 | 0.8762 |
| alder-straw | ko00910 | Nitrogen metabolism | 21 | 6 | 0.8466 |
| alder-straw | ko00780 | Biotin metabolism | 9 | 5 | 0.7978 |
| alder-straw | ko00740 | Riboflavin metabolism | 11 | 5 | 0.7602 |
| alder-straw | ko04978 | Mineral absorption | 7 | 4 | 0.727 |
| alder-straw | ko00380 | Tryptophan metabolism | 40 | 21 | 0.702 |
| alder-straw | ko05130 | Pathogenic Escherichia coli infection | 13 | 5 | 0.6742 |
| alder-straw | ko00640 | Propanoate metabolism | 28 | 13 | 0.6634 |
| solid-liquidExp | ko00970 | Aminoacyl-tRNA biosynthesis | 40 | 28 | 0.9998 |
| solid-liquidExp | ko01230 | Biosynthesis of amino acids | 116 | 58 | 0.9152 |
| solid-liquidExp | ko00520 | Amino sugar and nucleotide sugar metabolism | 49 | 28 | 0.8352 |
| solid-liquidExp | ko03030 | DNA replication | 32 | 27 | 0.7924 |

| | | | | | |
|---|---|---|---|---|---|
| solid-liquidExp | ko03008 | Ribosome biogenesis in eukaryotes | 64 | 31 | 0.7612 |
| solid-liquidExp | ko00280 | Valine, leucine and isoleucine degradation | 49 | 26 | 0.7048 |
| solid-liquidExp | ko04111 | Cell cycle - yeast | 75 | 43 | 0.6586 |
| solid-liquidExp | ko01524 | Platinum drug resistance | 29 | 18 | 0.6358 |
| solid-liquidExp | ko03050 | Proteasome | 35 | 18 | 0.6236 |
| solid-liquidSta | ko03008 | Ribosome biogenesis in eukaryotes | 64 | 38 | 1 |
| solid-liquidSta | ko00500 | Starch and sucrose metabolism | 65 | 38 | 0.997 |
| solid-liquidSta | ko00040 | Pentose and glucuronate interconversions | 35 | 25 | 0.9964 |
| solid-liquidSta | ko03030 | DNA replication | 32 | 22 | 0.983 |
| solid-liquidSta | ko00970 | Aminoacyl-tRNA biosynthesis | 40 | 22 | 0.9042 |
| solid-liquidSta | ko00052 | Galactose metabolism | 30 | 20 | 0.882 |
| solid-liquidSta | ko00564 | Glycerophospholipid metabolism | 39 | 22 | 0.8494 |
| solid-liquidSta | ko00770 | Pantothenate and CoA biosynthesis | 23 | 15 | 0.7834 |
| solid- | ko00965 | Betalain biosynthesis | 17 | 12 | 0.6102 |

| | | | | | |
|---|---|---|---|---|---|
| liquidSta | | | | | |
| stat-exp | ko00040 | Pentose and glucuronate interconversions | 35 | 21 | 1 |
| stat-exp | ko03010 | Ribosome | 99 | 29 | 1 |
| stat-exp | ko01120 | Microbial metabolism in diverse environments | 246 | 60 | 0.9772 |
| stat-exp | ko00280 | Valine, leucine and isoleucine degradation | 49 | 17 | 0.9676 |
| stat-exp | ko00520 | Amino sugar and nucleotide sugar metabolism | 49 | 12 | 0.8906 |
| stat-exp | ko00620 | Pyruvate metabolism | 41 | 11 | 0.8554 |
| stat-exp | ko00260 | Glycine, serine and threonine metabolism | 47 | 11 | 0.845 |
| stat-exp | ko00250 | Alanine, aspartate and glutamate metabolism | 31 | 9 | 0.7604 |
| stat-exp | ko01110 | Biosynthesis of secondary metabolites | 355 | 94 | 0.7382 |
| straw-malt | ko00500 | Starch and sucrose metabolism | 65 | 31 | 1 |
| straw-malt | ko00040 | Pentose and glucuronate interconversions | 35 | 19 | 1 |
| straw-malt | ko03008 | Ribosome biogenesis in eukaryotes | 64 | 23 | 1 |
| straw-malt | ko00330 | Arginine and proline metabolism | 38 | 18 | 0.9978 |
| straw-malt | ko00520 | Amino sugar and nucleotide sugar metabolism | 49 | 17 | 0.967 |
| straw-malt | ko00980 | Metabolism of xenobiotics by cytochrome P450 | 28 | 14 | 0.9512 |
| straw-malt | ko00630 | Glyoxylate and dicarboxylate metabolism | 40 | 21 | 0.9136 |
| straw-malt | ko00920 | Sulfur metabolism | 17 | 8 | 0.9056 |
| straw-malt | ko00052 | Galactose metabolism | 30 | 15 | 0.897 |
| straw-malt | ko00350 | Tyrosine metabolism | 50 | 20 | 0.838 |

| | | | | | |
|---|---|---|---|---|---|
| straw-malt | ko00770 | Pantothenate and CoA biosynthesis | 23 | 11 | 0.7994 |
| straw-malt | ko00910 | Nitrogen metabolism | 21 | 9 | 0.6518 |
| straw-malt | ko01220 | Degradation of aromatic compounds | 32 | 17 | 0.6144 |

#### Supplemental Table 3

| condition | GO ID | GO name | genes with this annotation in the genome | differentially expressed genes with this annotation | activation probability |
|---|---|---|---|---|---|
| alder-malt | GO:0016491 | oxidoreductase activity | 506 | 205 | 1 |
| alder-malt | GO:0005975 | carbohydrate metabolic process | 226 | 72 | 1 |
| alder-malt | GO:0071949 | FAD binding | 55 | 29 | 1 |
| alder-malt | GO:0055085 | transmembrane transport | 513 | 165 | 1 |
| alder-malt | GO:0055114 | oxidation-reduction process | 768 | 283 | 1 |
| alder-malt | GO:0008152 | metabolic process | 387 | 158 | 1 |
| alder-malt | GO:0008080 | N-acetyltransferase activity | 49 | 19 | 0.9998 |
| alder-malt | GO:0006508 | proteolysis | 132 | 39 | 0.7976 |
| alder-straw | GO:0016491 | oxidoreductase activity | 506 | 211 | 1 |
| alder-straw | GO:0003824 | catalytic activity | 610 | 212 | 1 |
| alder-straw | GO:0016787 | hydrolase activity | 165 | 57 | 1 |
| alder-straw | GO:0071949 | FAD binding | 55 | 28 | 1 |
| alder-straw | GO:0006508 | proteolysis | 132 | 53 | 1 |

| | | | | | |
|---|---|---|---|---|---|
| alder-straw | GO:0055114 | oxidation-reduction process | 768 | 301 | 1 |
| alder-straw | GO:0055085 | transmembrane transport | 513 | 176 | 0.9952 |
| alder-straw | GO:0005975 | carbohydrate metabolic process | 226 | 110 | 0.988 |
| solid-liquidExp | GO:0005524 | ATP binding | 497 | 231 | 1 |
| solid-liquidExp | GO:0005515 | protein binding | 749 | 314 | 1 |
| solid-liquidExp | GO:0003824 | catalytic activity | 610 | 256 | 1 |
| solid-liquidExp | GO:0055085 | transmembrane transport | 513 | 230 | 1 |
| solid-liquidExp | GO:0055114 | oxidation-reduction process | 768 | 341 | 0.9992 |
| solid-liquidExp | GO:0006508 | proteolysis | 132 | 68 | 0.9848 |
| solid-liquidExp | GO:0005975 | carbohydrate metabolic process | 226 | 115 | 0.9762 |
| solid-liquidExp | GO:0005634 | nucleus | 389 | 180 | 0.9716 |
| solid-liquidExp | GO:0016787 | hydrolase activity | 165 | 77 | 0.913 |
| solid-liquidExp | GO:0016491 | oxidoreductase activity | 506 | 231 | 0.8836 |
| solid-liquidExp | GO:0003676 | nucleic acid binding | 294 | 130 | 0.6472 |
| solid- | GO:0003824 | catalytic activity | 610 | 283 | 1 |

| | | | | | |
|---|---|---|---|---|---|
| liquidSta | | | | | |
| solid-liquidSta | GO:0005975 | carbohydrate metabolic process | 226 | 125 | 0.9898 |
| solid-liquidSta | GO:0055114 | oxidation-reduction process | 768 | 335 | 0.9868 |
| solid-liquidSta | GO:0008152 | metabolic process | 387 | 180 | 0.9798 |
| solid-liquidSta | GO:0055085 | transmembrane transport | 513 | 264 | 0.975 |
| solid-liquidSta | GO:0006508 | proteolysis | 132 | 66 | 0.8306 |
| stat-exp | GO:0003824 | catalytic activity | 610 | 164 | 1 |
| stat-exp | GO:0055114 | oxidation-reduction process | 768 | 232 | 1 |
| stat-exp | GO:0016491 | oxidoreductase activity | 506 | 153 | 0.9988 |
| stat-exp | GO:0071949 | FAD binding | 55 | 21 | 0.9874 |
| stat-exp | GO:0005975 | carbohydrate metabolic process | 226 | 67 | 0.9518 |
| stat-exp | GO:0055085 | transmembrane transport | 513 | 113 | 0.8948 |
| straw-malt | GO:0008080 | N-acetyltransferase activity | 49 | 22 | 1 |
| straw-malt | GO:0016491 | oxidoreductase activity | 506 | 192 | 1 |
| straw-malt | GO:0005975 | carbohydrate metabolic process | 226 | 117 | 1 |
| straw-malt | GO:0003824 | catalytic activity | 610 | 198 | 1 |
| straw-malt | GO:0016787 | hydrolase activity | 165 | 54 | 1 |

| | | | | | |
|---|---|---|---|---|---|
| straw-malt | GO:0071949 | FAD binding | 55 | 31 | 1 |
| straw-malt | GO:0006508 | proteolysis | 132 | 53 | 1 |
| straw-malt | GO:0055085 | transmembrane transport | 513 | 188 | 1 |
| straw-malt | GO:0055114 | oxidation-reduction process | 768 | 279 | 1 |
| straw-malt | GO:0000981 | RNA polymerase II transcription factor activity, sequence-specific DNA binding | 217 | 66 | 0.9982 |
| straw-malt | GO:0042254 | ribosome biogenesis | 15 | 9 | 0.634 |
